# No evidence for adaptation to local rhizobial mutualists in the legume *Medicago lupulina*

**DOI:** 10.1101/078675

**Authors:** Tia L. Harrison, Corlett W. Wood, Isabela L. Borges, John R. Stinchcombe

## Abstract

Local adaptation is a common but not ubiquitous feature of species interactions, and understanding the circumstances under which it evolves illuminates the factors that influence adaptive population divergence. Antagonistic species interactions dominate the local adaptation literature relative to mutualistic ones, preventing an overall assessment of adaptation within interspecific interactions. Here, we tested whether the legume *Medicago lupulina* is adapted to the locally abundant species of mutualistic nitrogen-fixing bacteria (“rhizobia”), which vary in frequency across its eastern North American range. We reciprocally inoculated northern and southern *M. lupulina* genotypes with the northern *(Ensifer medicae)* or southern bacterium *(E. meliloti)* in a greenhouse experiment. Neither northern nor southern plants produced more seed flowered earlier, or were more likely to flower when inoculated with their local rhizobium species, although plants produced more root nodules (the structures that house the bacteria) wit their local rhizobia. We used a pre-existing dataset to perform a genome scan for loci that showed elevated differentiation between field-collected plants that hosted different bacteria. None of the loci we identified belonged to the well-characterized suite of legume-rhizobia symbiosis genes, suggesting that the rhizobia do not drive genetic divergence between *M. lupulina* populations. Our results demonstrate that symbiont local adaptation is weak in this mutualism despite large-scale geographic variation in the identity of the interacting species.

## Introduction

Characterizing the circumstances under which local adaptation evolves informs our understanding of the relative importance of gene flow and selection, and thereby the extent and limitations of adaptive evolution (Antonovics, 1976; Bridle & Vines, 2007; Hereford, 2009; Savolainen *et al.,* 2013; Whitlock, 2015). However, existing tests of local adaptation to the biotic environment focus disproportionately on antagonistic interactions (but see Anderson et al. 2004, Hoeksema and Thompson 2007, Barrett et al. 2012), limiting our understanding of adaptation within the broad suite of interspecific interactions that occur in nature. Here we performed a reciprocal inoculation experiment to test for local adaptation in a classic mutualism: the symbiosis between legumes and nitrogen-fixing bacteria.

Local adaptation—when native genotypes outperform foreign genotypes in their home environment (Hereford 2009)—is driven by differences in selection in alternative environments, and is reflected in divergent phenotypes and genotypes between populations. The literature on local adaptation between interacting species is dominated by antagonistic species interactions such as those between hosts and their parasites, pathogens, or prey (Brodie *et al.,* 2002; Kawecki & Ebert, 2004; Hoeksema & Forde, 2008; Koskella *et al.,* 2012). Direct tests of symbiont local adaptation in mutualisms are rare (Hoeksema & Forde, 2008; Brockhurst & Koskella, 2013). Nevertheless, several lines of evidence suggest that adaptation to the local mutualist is a common feature of positive species interactions. Phenotype matching between local plant and pollinator communities is pervasive (Anderson *et al.,* 2009; Gómez *et al.,* 2009; Koski & Ashman, 2015), and a recent reciprocal translocation experiment showed that a plant’s reproductive success is highest in its local pollinator community (Newman *et al.,* 2015). In the classic mutualism between leguminous plants and nitrogen-fixing bacteria, genotype-by-genotype interactions— when fitness depends jointly on the genotypes of both partners—account for a substantial proportion of genetic variation in fitness-related traits within plant populations (Heath, 2010; Heath *et al.,* 2012; Ehinger *et al.,* 2014). On a broad geographic scale, these interactions are predicted to manifest as symbiont local adaptation when coupled with population differences in symbiont genotype frequencies (Heath & Nuismer, 2014).

Ultimately, though, directly testing for symbiont local adaptation in mutualisms requires assaying the fitness consequences of sympatric and allopatric symbionts in a reciprocal inoculation experiment (Heath & Stinchcombe, 2014). The diagnostic signature of symbiont local adaptation in these experiments is a genotype-by-genotype interaction for fitness, indicating that the fitness of one partner depends on the identity of its symbiont (Clausen *et al.,* 1940; Clausen & Hiesey, 1958; Kawecki & Ebert, 2004). Although these experiments are frequently used to test for parasite local adaptation in antagonistic interactions (reviewed in Hoeksema and Forde 2008), they are less commonly used to test for mutualist local adaptation (but see (Hoeksema & Thompson, 2007; Johnson *et al.,* 2010; Barrett *et al.,* 2012; Newman *et al.,* 2015)).

The economically and ecologically important mutualism between legumes in the genus *Medicago* and nitrogen-fixing bacteria (“rhizobia”) is well suited to testing for adaptation to the local mutualist (Cook *et al.,* 1997; Cook, 1999; Young *et al.,* 2011). In the facultative *Medicago-*rhizobia symbiosis, soil bacteria in the genus *Ensifer* (formerly *Sinorhizobium)* (Young, 2010) fix atmospheric nitrogen for their plant hosts in exchange for carbohydrates and housing in specialized root organs called nodules (Mylona *et al.,* 1995; van Rhijn & Vanderleyden, 1995). In eastern North America the relative frequencies of two principal symbionts *(Ensifer medicae* and *E. meliloti)* (Béna *et al.,* 2005) vary along a latitudinal cline (Figure S1) (Harrison *et al.,* 2017), which may generate strong selection on *Medicago* populations to adapt to their local *Ensifer* species. The bacteria are essential for plant growth in nitrogen-poor edaphic environments (Simonsen & Stinchcombe, 2014a), and genes mediating the association experience strong selection in both *Medicago* and *Ensifer* (Bailly *et al.*, 2006; De Mita *et al.*, 2007; Epstein *et al.,* 2012; Bonhomme *et al.,* 2015). Finally, there is substantial evidence for genotype-by-genotype interactions for fitness traits between *Medicago truncatula* and its *Ensifer* symbionts (Heath, 2010; Gorton *et al.,* 2012; Heath *et al.,* 2012), and suggestive evidence for some degree of co-speciation in the two genera (Béna *et al.,* 2005).

In the present study, we performed a reciprocal inoculation experiment to test for adaptation to the local rhizobia species in the black medic *(Medicago lupulina).* We tested the effect of sympatric and allopatric rhizobia on plant fitness in a greenhouse experiment. We measured traits closely associated with plant fitness (seed production) and rhizobial fitness (number of nodules formed with the plant host) to examine plant adaptation to their local rhizobial species and rhizobial adaptation to their local plant populations, respectively. Second, we took advantage of an existing genomic data set and performed a genome scan for loci that exhibited elevated differentiation between field-collected plants associated with different bacterial species in natural populations. Genome scans identify loci that exhibit heightened differentiation between populations inhabiting alternative environments, which are presumed to constitute the genetic basis of local adaptation (Coop *et al.,* 2010; Günther & Coop, 2013; Savolainen *et al.,* 2013; Tiffin & Ross-Ibarra, 2014). Unlike reciprocal inoculation experiments, these tests integrate across generations and ancillary environmental variation, capturing the cumulative effects of long-term selection in alternative environments (Tiffin & Ross-Ibarra, 2014; de Villemereuil *et al.,* 2015; Jensen *et al.,* 2016).

Neither the phenotypic nor genomic approaches revealed strong evidence of adaptation to the local rhizobia in *M. lupulina.* We found suggestive evidence that the rhizobia may be adapted to their local plant populations; plants formed more nodules with rhizobia from the same geographic region. However, overall our data suggest that symbiont local adaptation appears to be weak or absent in this mutualism’s North American range.

## Materials and Methods

### Study system

*Medicago lupulina* is an annual, highly self-fertilizing legume native to Eurasia (Turkington & Cavers, 1979; Yan *et al.,* 2009). After its introduction to North America in the 1700s, *M. lupulina* expanded its range to occupy nitrogen-poor areas of the continent’s temperate and subtropical regions (Turkington & Cavers, 1979). *Medicago* has a short generation time (Turkington & Cavers, 1979), its rhizobia are easily manipulated (Heath & Tiffin, 2007), an annotated genome is available in the genus (Young *et al.,* 2011), and the genes involved in the rhizobial mutualism are extensively characterized (Mylona *et al.,* 1995; Cook *et al.,* 1997; Young *et al.,* 2011).

The relative frequencies of *M. lupulina’s* two symbiotic rhizobia species *(Ensifer medicae* and *E. meliloti)* vary along a northwest-to-southeast cline in eastern North America (Figure S1) (Harrison *et al.,* 2017). Moreover, there is very little genetic diversity within each *Ensifer* species at neutral sites and at known symbiosis genes (Harrison *et al.,* 2017). Whole-genome sequencing of *Ensifer* strains sampled from the same populations used in the present study found that neutral diversity extremely low in *E. medicae* (π_synonymous_ = 0.0006 and in *E. meliloti* (π_synonymous_ = 0.0001). Crucially, there was also very little diversity in genes known to be involved in the legume-rhizobia symbiosis. There were few or no SNPs in any nodulation genes (nodA, -B, or - C), nitrogen fixation genes (nifA, -B, -D, -E, -H, -N, or -X), or type III effector genes in either *E. medicae* (π ≤ 0.0004 for these genes) or *E. meliloti* (π ≤ 0.0002). Taken together, this evidence suggests that there is very little functional diversity within *Ensifer* species in North America, so we only tested a single strain per species in the reciprocal inoculation experiment described below.

### Reciprocal inoculation experiment

To test for adaptation to the local rhizobia, we inoculated *M. lupulina* genotypes from the northern and southern portions of the plant’s eastern North American range with either the locally abundant rhizobium species in the north *(E. medicae)* or in the south *(E. meliloti).* From a total of 39 *M. lupulina* populations sampled between Delaware and Ontario in September-October of 2013 (Harrison *et al.,* 2017), we selected 7 southern and 7 northern plant populations in which Harrison et al. (2017) detected only a single *Ensifer* species (Figure 2, Table S1; see Figure S1 for a complete map with all 39 sampled populations). Within each population, seeds and root nodules were collected from 2-10 randomly chosen *M. lupulina* individuals. All sampled plants were at least 0.5m apart. Nodules were stored at 4°C in plastic bags until they were processed. Field-collected seeds from these populations were grown in the greenhouse for one generation to reduce maternal and environmental effects from the field, and we performed our experiments using the progeny of these greenhouse-grown plants.

**Figure 1.**
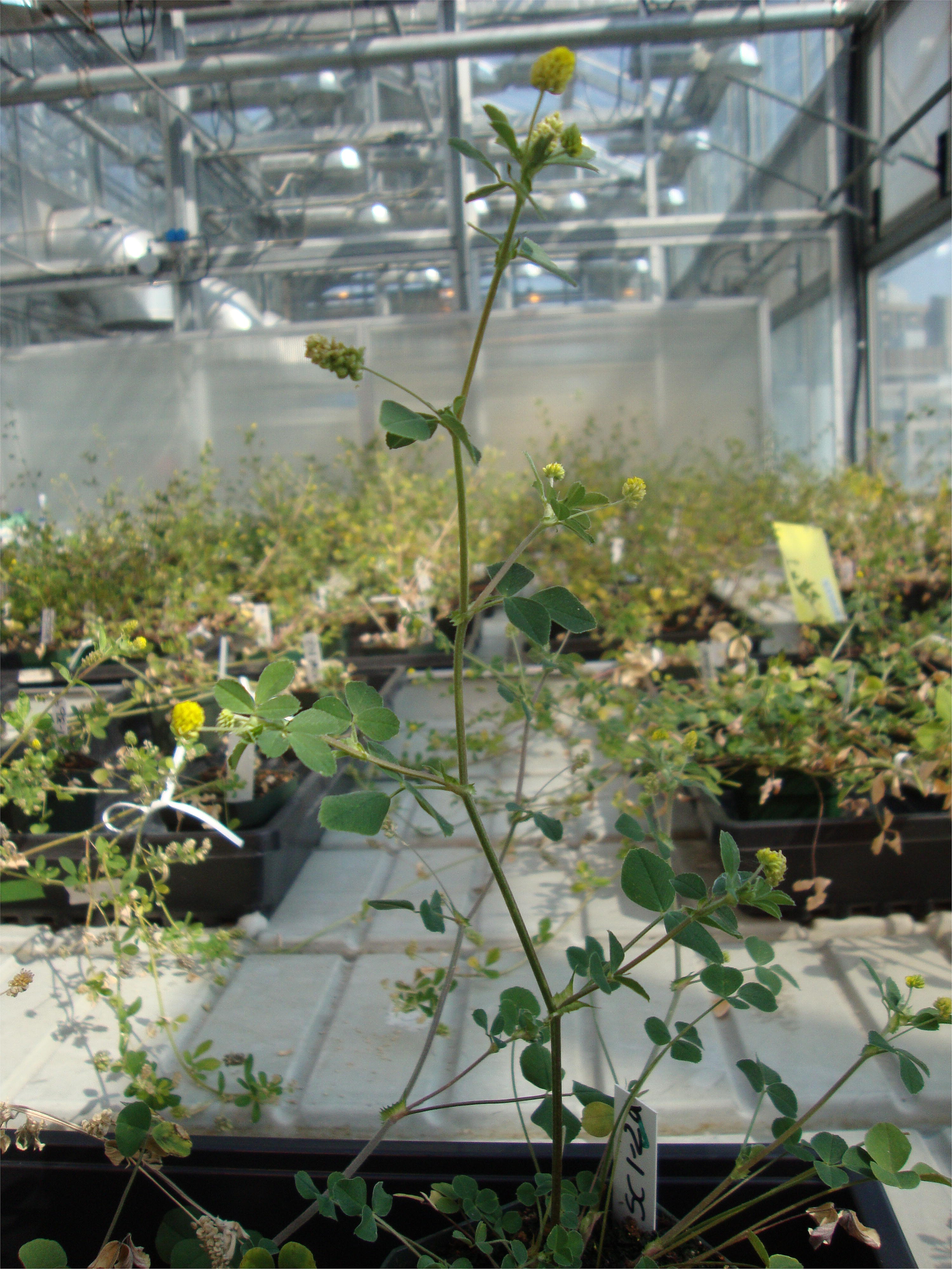
A *Medicago lupulina* individual flowering in the greenhouse.

**Figure 2.**
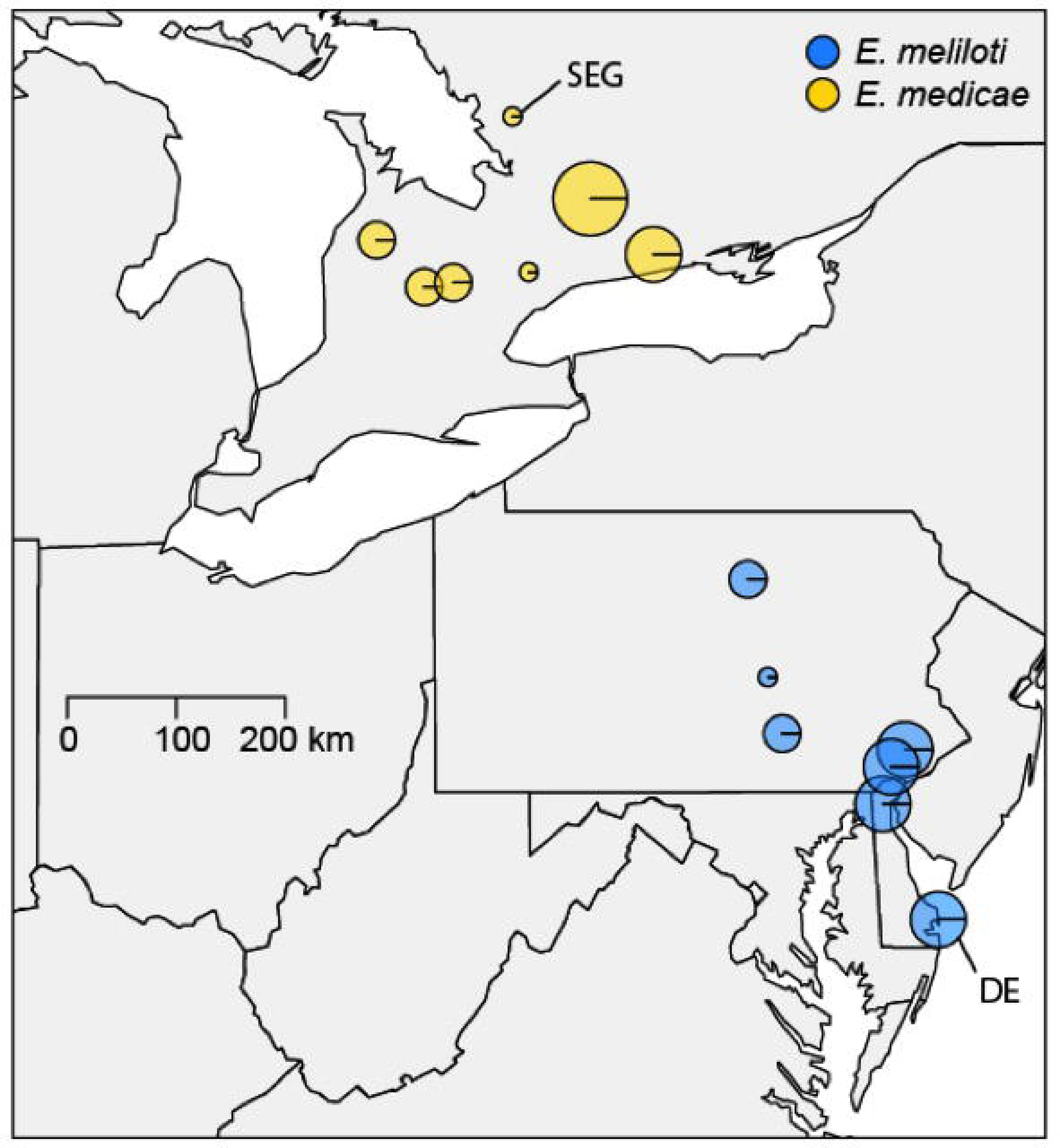
Locations of the 14 *M. lupulina* populations used in this study. The size of each circle corresponds to the number of plants sampled from the population, and the color indicates the rhizobia. The *E. medicae* strain used in the reciprocal inoculation experiment was obtained from the northernmost population sampled (“SEG”); the *E. meliloti* strain was obtained from the southernmost population (“DE”). See Table S1 for GPS coordinates.

We planted F_1_ greenhouse-derived seeds of 43 maternal families (27 from the north and 16 from the south) in a split-plot randomized complete block design in the greenhouse at the University of Toronto. Seedlings were planted with sterile forceps into cone-tainers filled with sand (autoclaved twice at 121°C). *Medicago lupulina* naturally colonizes sandy soils (Turkington and Cavers 1979), and the plants rely heavily on nitrogen supplied by their rhizobial symbionts under these conditions because sand is a nitrogen-poor substrate (Heath et al. 2010). Each block was divided into two bacterial treatments, each containing 15 northern and 11 southern plants, the locations of which were randomized within blocks. Populations were split across blocks. Due to seed limitations, not all families were represented in every block, but within a block both bacterial treatments comprised the same 26 families. We replicated this design across six blocks, for a total of 312 plants (6-13 replicates per family for 37 families; 1-4 replicates per family for 6 families). An additional block containing 42 plants (33 from the north and 9 from the south) served as an inoculation control, and a means for estimating plant performance and fitness in the absence of either bacterial species. Prior to planting, seeds were scarified with a razor blade, sterilized with ethanol and bleach, and stratified on 8% water agar plates at 4°C for 7 days to germinate. We misted seedlings with water daily and fertilized with 5mL of nitrogen-free Fahraeus medium (noble.org/medicagohandbook) twice before inoculation with rhizobia.

The *Ensifer* strains used for inoculation were recovered from frozen samples collected by Harrison et al. (2017) from two of the populations used in our experiment. The strains were originally cultured from field-collected root nodules by sterilizing one nodule per plant in ethanol and bleach, and crushing and plating it onto a 2% tryptone yeast (TY) agar plate. Strains were re-streaked onto TY agar four times to reduce contamination and grown at 30°C for 48 hours, after which they were transferred to liquid TY media and cultured for two days at 30°C. To identify each strain to species *(E. medicae* or *E. meliloti),* DNA was extracted from liquid cultures (cell density: 8 x 10^8^ cells/ml) using the MoBio UltraClean Microbial DNA Isolation Kit, whole-genome sequenced at SickKids Hospital (Toronto, Ontario), and genotyped using GATK (McKenna *et al.,* 2010). We used alignment scores and the *Ensifer* 16S locus (Rome *et al.,* 1997) to determine species identity of rhizobia strains associated with the sampled plants.

We selected one *E. medicae* strain from the northernmost population in Ontario and one *E. meliloti* strain from the southernmost population in Delaware for our experiment (“SEG” and “DE” in Figure 2). Genetic diversity is very low among strains within *Ensifer* species across North America (see "Study system" above) (Harrison *et al.,* 2017), so the specific strains used are not likely to influence our results. Prior to inoculation, these strains were cultured as described above from samples stored at −80°C. Liquid cultures were diluted with sterile TY media to an OD600 reading of 0.1 (a concentration of ~10^6^ cells per mL) (Simonsen & Stinchcombe, 2014b). Each plant received 1 mL of inoculate 13 days after planting, and 1 mL again 10 days later. Controls were also inoculated twice with sterile TY media 10 days apart, and were used to assess rhizobia contamination across treatments. Throughout the remainder of the experiment, all plants were bottom-watered three times a week. We used two bottom-watering trays per block, such that all plants in a given bacterial treatment had the same tray, while those from the alternative bacterial treatment had a different tray.

We scored mortality weekly throughout the experiment, counted the number of leaves every 4 weeks, recorded the date of first flower, and collected seeds. After five months, which approximates the length of the April-October growing season in southern Ontario (Turkington & Cavers, 1979), we harvested all plants and collected any remaining unripe seeds. We dried and weighed aboveground tissue from each plant to the nearest 0.1 mg, and counted all seeds and root nodules (symbiotic organs housing the rhizobia).

We analyzed five traits to test whether northern and southern *M. lupulina* plants were adapted to their local rhizobium: number of seeds, aboveground biomass, flowering time (excluding plants that did not flower), probability of flowering, and number of nodules. All analyzes were performed in R v.3.2.4 with sum-to-zero contrasts (“contr.sum”) (R Core Team, 2016), and we tested significance using type III sums of squares in the function Anova in the *car* package (Fox & Weisberg, 2011). Log-transformed aboveground biomass and flowering time were analyzed with general linear mixed models using the function lmer in the *lme4* package (Bates *et al.,* 2015). Probability of flowering and number of nodules were analyzed with generalized linear mixed models with binomial and Poisson error distributions, respectively, using the function glmer in the *lme4* package (Bates *et al.,* 2015). We verified that all dependent variables met the assumptions of linearity, normality, and homoscedasticity through visual inspection of quantile-quantile plots, plots of the residuals versus fitted values, and scale-location plots. Seed number was severely zero-inflated (42% of plants did not produce seeds), so we analyzed it using a mixture model (see below).

Each of the above models included rhizobia treatment *(E. medicae* or *E. meliloti),* region (north or south), and the rhizobia-by-region interaction as fixed effects. A significant rhizobia-by-region interaction, in which northern plants have higher fitness when inoculated with *E. medicae* and southern plants have higher fitness with *E. meliloti,* would be evidence for symbiont local adaptation. We included a fixed effect of researcher in our analysis of nodule counts.

Block, population, and family nested within population were included as random effects. We also included the block-by-treatment interaction as a random effect because the rhizobia treatment was applied at the half-block rather than at the plant level (Altman & Krzywinski, 2015). While this design provides a weaker test of the rhizobia main effect, it is sensitive to the detection of rhizobia-by-region interactions, the main goal of our experiment (Altman & Krzywinski, 2015). We tested the significance of the population and family random effects in our analyses of biomass, number of nodules, and flowering time using likelihood ratio tests in which we compare the full model to reduced models without the random effects. Because these likelihood ratio tests test a null hypothesis located at the boundary of parameter space (i.e., a variance of zero), the p-values produced by these tests are twice what they should be, so we divided these p-values in half (Bolker *et al.,* 2009).

We analyzed seed number with a zero-inflated Poisson model implemented with the function MCMCglmm in the package *MCMCglmm* (Hadfield, 2010). Zero-inflated models are a type of mixture model in which the zero class is modeled as the combined result of binomial and count processes (Zuur *et al.,* 2009). In MCMCglmm, zero-inflated Poisson GLMMs are fit as multi-response models with one latent variable for the binomial zero-generating process and one for the Poisson count-generating process (Hadfield, 2015). We fit a model for seed number that included fixed effects of rhizobia, region, the rhizobia-by-region interaction, and the reserved MCMCglmm variable "trait" that indexes the binomial and Poisson latent variables. We omitted the interaction between trait and other fixed effects in order to estimate a single effect of rhizobia, region, and the rhizobia-by-region interaction across both the binomial and Poisson processes. Block, population, family, and the block-by-treatment effect were included as random effects. Different random effect variances were fit to the binomial and Poisson processes using the "idh" variance structure in MCMCglmm (Hadfield, 2015). We fit a residual variance (R) structure using the argument rcov = ~ us(trait):units, which allows a unique residual for all predictors in the model, used the default priors for the fixed effects (mean = 0, variance = 10^10^) and specified parameter-expanded priors (alpha.mu = 0, alpha.v = 1000) for the random effects (Hadfield, 2010).

We ran the model for 1,300,000 iterations, discarded the first 300,000 iterations, and stored every 1,000^th^ iterate. Model convergence was assessed with traceplots, running mean plots, and autocorrelation plots of the fixed and random effects using the *coda* (Plummer *et al.,* 2006) and *mcmcplots* (McKay Curtis, 2015) packages. Even though we used parameter-expanded priors on the random effects, the estimates of the population and block random effects remained close to zero, but omitting these terms from our model did not qualitatively change the results. We assessed significance of the population and family random effects in our analysis of seed number by comparing the Deviance Information Criteria (DIC; a Bayesian analog of AIC) of the full model to reduced models that omitted the random effects (Spiegelhalter *et al.,* 2002; Bolker *et al.,* 2009). We considered a random effects to be significant when the reduced model had ΔDIC greater than two, indicating that it was a worse fit to the data than the full model (Spiegelhalter *et al.,* 2002).

Finally, we calculated pairwise correlations between all traits using Spearman’s correlation on the family means for each trait. We obtained family means for biomass, flowering time, and number of nodules by extracting the conditional modes (also known as the best linear unbiased predictors, or BLUPs) for each level of the family random effect from the models described above. For number of seeds, we used the marginal posterior modes of the family random effect as our family mean estimates.

### Genomic data set

A limitation of using reciprocal inoculation experiments to test for symbiont local adaptation is that the fitness benefit of a symbiosis often depends on the biotic and abiotic environmental conditions in which it is expressed (Heath & Tiffin, 2007; Heath *et al.,* 2010; Porter *et al.,* 2011; Barrett *et al.,* 2012; Simonsen & Stinchcombe, 2014a). To address this limitation, we took advantage of a pre-existing *M. lupulina* SNP dataset collected by Harrison et al. (2017) to perform genomic scans in *M. lupulina.* The goal of this analysis was to determine whether genes involved in the legume-rhizobia symbiosis are differentiated between plants associated with different *Ensifer* species in natural populations, a pattern that would be consistent with symbiont local adaptation.

Details on SNP discovery methods can be found in Supplemental Methods (Appendix 1), Harrison et al. (2017). In brief, field-collected seeds from 73 *M. lupulina* individuals were grown in the greenhouse as described in the "Reciprocal inoculation experiment" section above. We extracted DNA from leaf tissue collected from one individual per maternal line and samples were sequenced at Cornell University using genotyping-by-sequencing (GBS) in two Illumina flow cell lanes (Elshire *et al.,* 2011). Genomic libraries were prepared with the restriction enzyme EcoT22I, and SNPs were called using the program Stacks (Catchen *et al.,* 2011, 2013). We extracted and sequenced rhizobia DNA from one nodule from each field-sampled plant, and determined the species identity of each strain as described in the "Reciprocal inoculation experiment" section above.

We searched for outlier loci between *M. lupulina* plants hosting *E. medicae* and *E. meliloti* to assess whether there is evidence for genetic divergence between plants associated with different *Ensifer* species. We used the program Bayenv2 to calculate X^T^X statistics for each SNP in the *M. lupulina* sample (Coop *et al.,* 2010): X^T^X is an F_ST_-like statistic that controls for population variation and covariation in allele frequencies (i.e., population structure). We estimated the covariance matrix using 100,000 iterations. Because we only wanted to calculate X^T^X statistics and did not wish to calculate environmental correlations, we included an environmental file of dummy values to run Bayenv2 but avoid environmental analysis. We ranked SNPs from highest to lowest X^T^X values and identified the top 1% of SNPs to BLAST against the reference genome of *M. truncatula* to identify the outlier loci involved in rhizobia association in *M. truncatula* (taxonomy ID 3880) (Tang *et al.,* 2014). We aligned to the *M. truncatula* genome to leverage its extensive genomic resources; *M. truncatula* is the closest relative of *M. lupulina* with a thoroughly mapped and annotated genome. We used nucleotide BLAST (blastn) to search somewhat similar sequences in the unannotated *M. truncatula* genome in order to retrieve chromosome positions for our outlier loci. To identify the orthologous gene associated with each outlier locus, we then looked up the chromosome position of each outlier in the annotated *Medicago truncatula* genome (Mt. 4.0 http://jcvi.org/medicago/). We performed the outlier loci test in three ways. First, we characterized outlier loci using the range-wide sample of plants (73 plant individuals) and compared the results to outlier loci found using the southern Ontario samples (49 plant individuals) to account for possible covariance between environmental gradients and bacterial species composition. Second, because outlier loci are usually in linkage disequilibrium (LD) with the causal genes responsible for adaptation, we searched for legume-rhizobia symbiosis genes within 5 kb and 10 kb of our detected outlier loci (Branca *et al.* 2011). Third, we measured the distance in base pairs between the *M. truncatula* orthologs of detected outlier loci and key *M. truncatula* genes involved in the rhizobia symbiosis, assuming synteny between *M. lupulina* and *M. truncatula.* Details of these analyses can be found in the Supplemental Methods (Appendix 2).

## Results

### Reciprocal inoculation experiment

Uninoculated *Medicago lupulina* plants performed extremely poorly without rhizobia. None of our uninoculated control plants flowered or set seed, and the biomass of control plants was approximately 20-fold smaller than inoculated plants (least squares mean ± SE (mg): controls: 21.01 ± 0.05; inoculated plants from both rhizobia treatments: 476.01 ± 0.03; F_1,14.808_ = 610.7, P < 0.001). The performance of the control plants also demonstrates that cross-contamination between the two rhizobia treatments was likely minimal in our experiment. Only 1 of 42 uninoculated control plants produced nodules, and this anomalous individual was similar in size to the rest of the controls for the first several months, indicating that it probably did not nodulate until late in the experiment.

In plants inoculated with *E. medicae* or *E. meliloti,* pairwise family mean correlations between all measured traits were generally low, indicating that the traits that we measured were largely independent of one another (r ≤ |0.10|, P ≥ 0.54). Only flowering time and aboveground biomass were significantly correlated (r = 0.49, P = 0.002); later-flowering plants had greater aboveground biomass.

Our analysis of seed number, probability of flowering, and flowering time revealed no evidence of adaptation to the local rhizobia. There was no significant rhizobia-by-region interaction for any of these reproductive traits (Figure 3, Table 1). There was a marginally significant effect of region on seed number; southern plants produced more seeds than northern plants in both rhizobia treatments (Figure 3A, Table 1). There was no significant effect of rhizobia treatment or region on either flowering trait (Figure 3C, Table 1).

**Figure 3.**
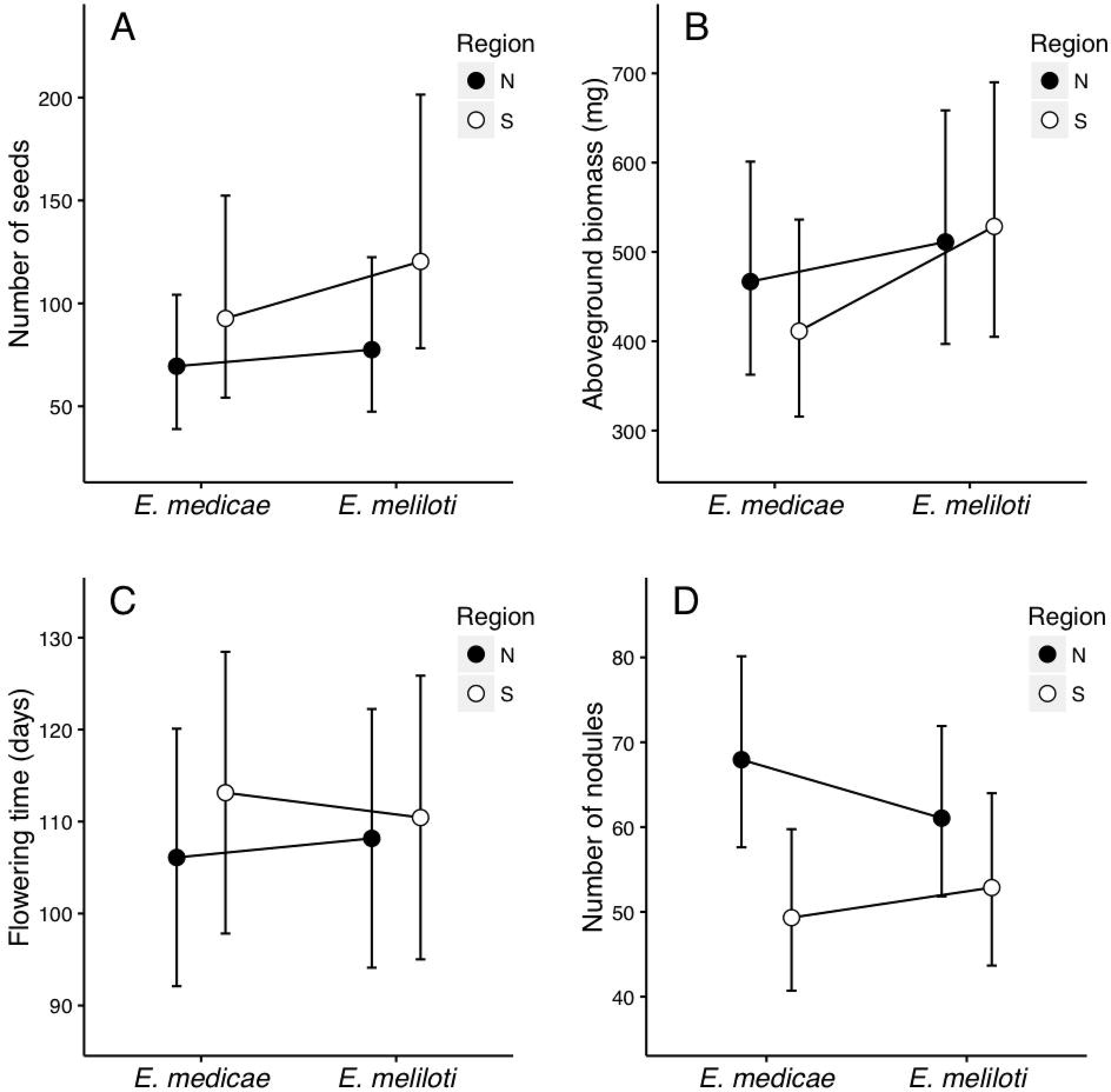
Least squares means and 95% confidence intervals for northern (black) and southern (white) plants grown in the two rhizobia treatments. *Ensifer medicae* is the locally abundant rhizobia in the north, and *E. meliloti* is the locally abundant rhizobia in the south. (A) Number of seeds; (B) aboveground biomass; (C) flowering time; (D) number of nodules.

**Table 1.**
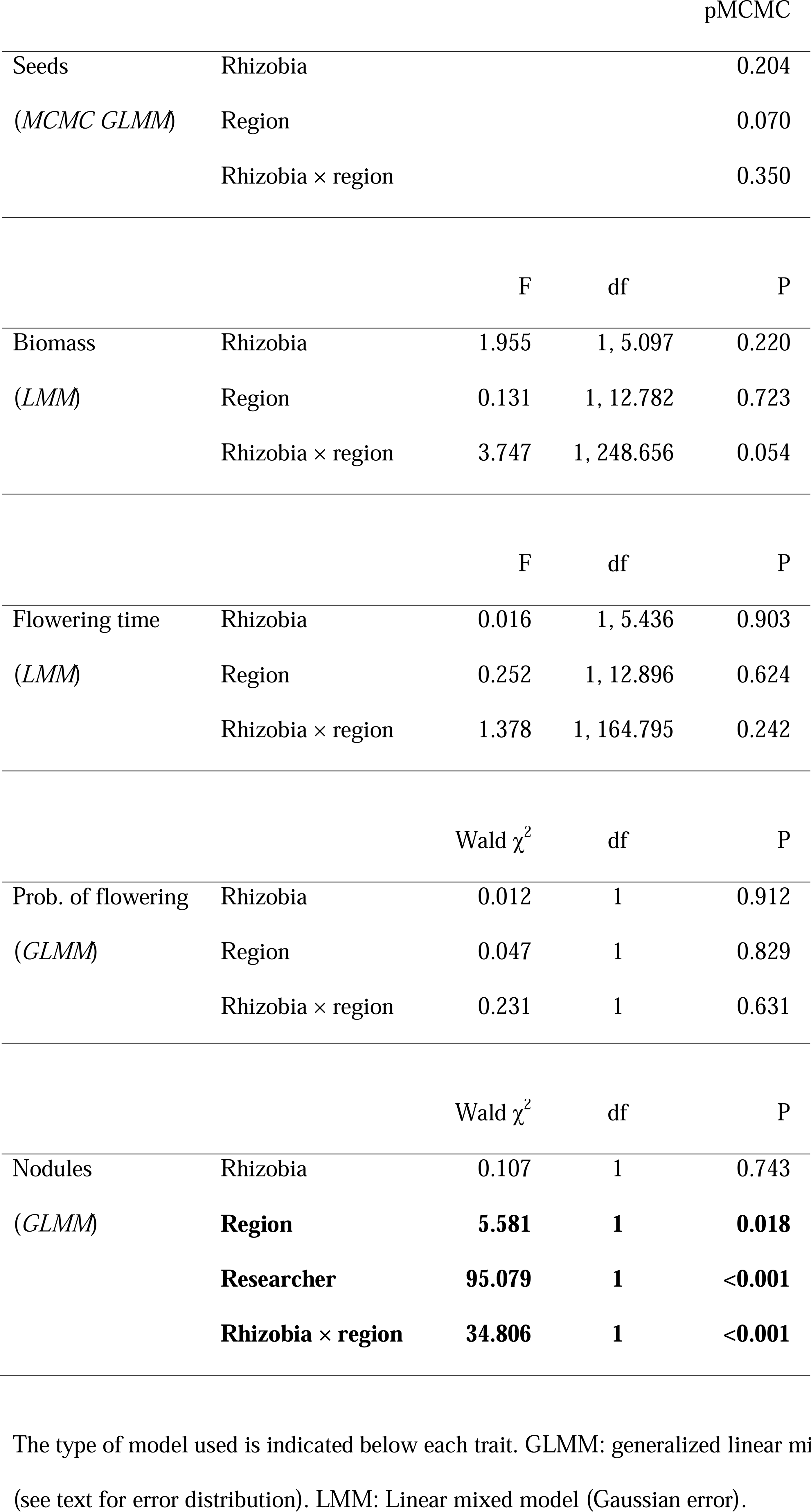
Results of general(ized) linear mixed models testing for local adaptation in the reciprocal inoculation experiment.

The rhizobia-by-region interaction for aboveground biomass was marginally significant (P_rhizobia-by-region interaction_ = 0.054, Table 1). While the biomass of northern plants was unaffected by rhizobia treatment, southern plants produced more aboveground biomass when inoculated with *E. meliloti* (Figure 3B), the locally abundant rhizobia in south.

We found a highly significant rhizobia-by-region interaction for nodule number (Table 1). Northern plants produced more nodules that southern plants when inoculated with *E. medicae,* the locally abundant rhizobia in the north. The difference between northern and southern plants decreased when inoculated with *E. meliloti,* an effect that was driven by both an increase in nodulation in southern plants and a decrease in nodulation in northern plants (Figure 3D). There was also a significant effect of region, indicating that northern plants produced more nodules across both rhizobia treatments, and a significant effect of researcher (Table 1).

Aboveground biomass varied significantly among populations (χ^2^ = 3.405, P = 0.033) and families (χ^2^ = 40.490, P < 0.001). Number of nodules did not vary among populations (χ^2^ = 0, P = 0.500), but did vary among families (χ^2^ = 1305.200, P <0.001). Flowering time varied significantly among populations (χ^2^ = 12.058, P <0.001) and families (χ^2^ = 28.511, P <0.001). Number of seeds did not vary significantly among populations (ΔDIC_reduced_ = 1.54), but did vary among families (ΔDIC_reduced_ = 133.7).

### Genomic outlier analysis

We identified three outlier loci that appeared in the top 1% of SNPs in both the rangewide *M. lupulina* sample and southern *M. lupulina* Ontario sample in our Bayenv2 analysis (Supplemental Table S2 and S3). None of these three loci mapped to a specific gene in the *M. truncatula* reference genome. Furthermore, we did not find any genes involved in the legume-rhizobia interaction within 5 kb or 10 kb of our three outlier loci (Supplemental Table S4). Finally, the base pair distances between our outlier loci and known genes involved in the *Medicago-Ensifer* mutualism were very large (minimum: 18 kb) (Supplemental Table S4).

Details on summary X^T^X statistics, BLAST alignment scores, and gene functions are presented in the Supplemental Methods (Appendix 3) and Supplemental Table S2.

## Discussion

We performed a reciprocal inoculation experiment and genome scan to test for symbiont local adaptation in the mutualism between *M. lupulina* and nitrogen-fixing *Ensifer* bacteria across its eastern North American range. We found no evidence that plants are adapted to the locally abundant rhizobia species in either analysis. Local rhizobia did not have differential fitness consequences for their host plants, nor were any of the well-characterized symbiosis genes differentiated between field-collected plants associated with different rhizobia. However, we did find suggestive evidence for rhizobium local adaptation to the plant host; plants produced more nodules with rhizobia from the same geographic region. Overall, symbiont local adaptation appears to be absent or weak in this mutualism’s eastern North American range despite the strong cline in the relative abundances of the two rhizobia species.

### Reciprocal inoculation experiment & genomic outlier analysis

Uninoculated plants performed extremely poorly without either *Ensifer* species, demonstrating that *M. lupulina* is adapted to symbiosis with rhizobia. Despite differential nodulation with local and foreign rhizobia (P_rhizobia-by-region_ < 0.001, Table 1), however, there was no strong evidence for adaptation to the local rhizobia in other plant traits. One explanation for this pattern is that plants modify their nodulation strategy to compensate for differences in symbiotic efficiency with local and foreign rhizobia. The congeneric species *M. truncatula* adjusts its nodulation strategy in response to the rhizobia nitrogen fixation efficiency (Heath & Tiffin 2009), which jointly depends on plant and rhizobia genotype (Mhadhbi *et al.,* 2005). If plants produce more nodules with less efficient symbionts, increased nodulation may not translate to greater nitrogen uptake, masking any effects of differential nodulation on biomass and seed production. The fact that seed number, a reasonable proxy for total fitness in a selfing annual or short-lived perennial like *M. lupulina* (Turkington & Cavers, 1979), was unaffected by the local rhizobia strongly suggests that adaptation to the local rhizobia was absent in our experiment at the whole-plant level.

Our data suggest that symbiont local adaptation may be restricted to the rhizobia in this mutualism; the rhizobia may be adapted to their local *M. lupulina* genotype even though the plant does not appear to be adapted to its local rhizobium. The strongest signature of local adaptation in our reciprocal inoculation experiment occurred in nodule traits, a pattern that has also been documented in congeneric *Medicago* species (Porter *et al.,* 2011). Differential nodulation may impact the rhizobia more than the plant, given that nodule number is correlated with rhizobia fitness in *Medicago* (Heath, 2010).

However, even in the nodulation trait that exhibited a strong rhizobia-by-region interaction—the statistical signature of local adaptation—the data are only weakly consistent with the canonical pattern of local adaptation. The strongest test of local adaptation is whether local genotypes outperform foreign genotypes in all environments (the "local-versus-foreign" criterion) (Kawecki & Ebert, 2004). Neither trait that exhibited any rhizobia-by-region interaction (number of nodules and aboveground biomass) satisfied this criterion. Instead, our results were more closely aligned with a weaker test of local adaptation, which diagnoses local adaptation when each genotype’s fitness is greater in its native environment than in alternative environments (the "home-versus-away" criterion) (Kawecki & Ebert, 2004).

Although reciprocal inoculation experiments are powerful because they reflect whole-organism performance in native and foreign environments, genotype-by-environment interactions are sensitive to experimental conditions (Kawecki & Ebert, 2004). We designed our experiment to minimize this concern by mimicking key abiotic conditions relevant to the *Medicago-rhizobia* mutualism. For example, we planted in sand, which reflects the nitrogen-poor stony roadside soils that *M. lupulina* naturally colonizes (Turkington and Cavers 1979). Under such nitrogen-limited conditions, the mutualism between legumes and nitrogen-fixing bacteria is especially crucial for the plant (Heath et al. 2010). Nevertheless, the absence of local adaptation in our experiment could be due to experimental conditions not adequately reflecting the typical natural environment (in our case, cone-tainers, sterilized greenhouse soil, artificial day length control, absence of other biotic interactors, etc.). A second caveat of our reciprocal inoculation experiment is that we used only one strain for each rhizobium species. Although previous work suggests that functional diversity within *E. medicae* and *E. meliloti* is extremely limited in eastern North America, it is possible that we would have detected a signal of local adaptation had we used different or larger sample of rhizobia strains.

To address these caveats, we performed a genomic outlier analysis to search for allele frequency differences between plants associated with different rhizobia species. Genome scans circumvent the caveats described above because they integrate across many generations of selection and ancillary environmental variation. However, we also found very weak evidence of symbiont local adaptation in this analysis. The loci that were highly differentiated between plants hosting different *Ensifer* species (the top 1% of loci in the X^T^X outlier analysis) were not associated with any genes involved in the legume-rhizobia symbiosis in either the range-wide or Ontario samples. Moreover, none of the *M. truncatula* orthologs of our outlier loci were located within the scale of linkage disequilibrium (5-10 kb in *M. truncatula)* (Branca *et al.,* 2011) from known symbiosis genes. It is unlikely that the loci identified in our genome scan are novel *M.* lupulina-specific symbiosis genes underlying adaptation to the local bacteria. The *Medicago* genes involved in symbiotic interactions with rhizobia are well characterized and highly conserved in legumes (Rostas *et al.,* 1986; van Rhijn & Vanderleyden, 1995; De Mita *et al.,* 2006; Branca *et al.,* 2011; Gorton *et al.,* 2012; Stanton-Geddes *et al.,* 2013). *Medicago lupulina* is a close relative of *M. truncatula* (Bena, 2001; Yoder *et al.,* 2013), and both plants fix nitrogen with both *Ensifer* species tested in our experiment (Béna *et al.,* 2005). However, our results are subject to caveats common to genome scans for selection (Pavlidis *et al.,* 2012). In particular, our sample size in terms of individuals, and the number of SNPs, was low, reducing our power. Therefore, our genome scan might not have been able to detect highly differentiated loci important for symbiont local adaptation in *M. lupulina*.

### Local adaptation in the legume-rhizobia symbiosis

Our data suggest that symbiont local adaptation may be asymmetric in this mutualism. Although plant fitness was not affected by the identity of its rhizobial symbiont, plants tended to produce more nodules with rhizobia from the same geographic region. This pattern was especially pronounced for the northern rhizobium, *E. medicae.* Stronger symbiont local adaptation in one partner commonly occurs in host-parasite systems (Hoeksema & Forde, 2008), but the phenomenon has not been systematically explored in the context of mutualism even though asymmetrical evolutionary rates in coevolving species pairs are expected in both mutualisms and antagonisms (Bergstrom & Lachmann, 2003).

Nevertheless, our phenotypic and genomic data indicate that *M. lupulina* is not adapted to the local rhizobia across its eastern North American range. The absence of symbiont local adaptation in this mutualism is surprising given that the system is characterized by several features that ordinarily strongly favor its evolution. Genotype-by-genotype interactions commonly occur between a congener, *M. truncatula,* and different strains of the same *Ensifer* species (Heath & Tiffin, 2007; Heath, 2010; Heath *et al.,* 2012), suggesting that the genetically divergent rhizobia *species* (Bailly *et al.,* 2006) we assayed would have even greater effects on their plant host. Furthermore, there is a cline in the frequencies of the two rhizobia across a large geographic scale that coincides with plant population genetic structure (Harrison *et al.,* 2017). What might account for the lack of symbiont local adaptation in this mutualism?

Gene flow may overwhelm the effects of local selection, leading to a low equilibrium level of genetic differentiation between plants associated with different rhizobia (McKay & Latta, 2002). Although there is a strong geographic cline in the frequencies of the two *Ensifer* species, Harrison et al. (2017) did detect *E. meliloti* in some northern populations and *E. medicae* in some southern populations. Symbiont local adaptation within *M. lupulina* populations could be swamped by gene flow from neighboring populations that encounter the alternative mutualist, or by the invasion of the alternative mutualist itself. Frequent gene flow across large geographic distances is consistent with the ecology of *M. lupulina* in North America. It is dispersed over long distances by birds, grazing animals, and water, and is likely distributed as a component in seed mixes for forage crops as well (Turkington and Cavers 1979).

Gene flow in the form of horizontal gene transfer between the two rhizobia species could similarly homogenize any signature of local selection (Lenormand, 2002; Bailly *et al.,* 2007). Bacteria that form nitrogen-fixing symbioses with legumes have been shown to horizontally transfer genes involved in forming and maintaining the mutualism (Suominen *et al.,* 2001; Aoki *et al.,* 2013; Lemaire *et al.,* 2015). Swapping cassettes of symbiosis genes would enable genetically divergent rhizobial species to associate with the same hosts, largely eliminating among-symbiont differences from the perspective of the plant. Finally, temporal variation in the biotic and abiotic environment may modify the costs and benefits of the mutualism (Heath *et al.,* 2010; Heath & McGhee, 2012; Simonsen & Stinchcombe, 2014a), weakening selection favoring local rhizobia.

Alternatively, symbiont local adaptation may generate relatively weak fitness tradeoffs in mutualisms. The fitness tradeoffs that are the hallmark of local adaptation evolve whenever adaptation to one environment results in maladaptation to another (Kawecki & Ebert, 2004). It has been hypothesized that selection in coevolving mutualisms strongly favors general compatibility and the reduction of fitness tradeoffs (Law & Koptur, 1986; Parker, 1999; Barrett *et al.,* 2012). Selection to minimize fitness tradeoffs may be especially strong in the legume-rhizobia mutualism, which is crucial for plants growing in nitrogen-poor soils (Heath *et al.,* 2010). Under nitrogen-limited conditions, the cost of maladaptation to a locally rare rhizobium may be severe enough to outweigh the selective advantage of a marginal increase in the benefits obtained from the locally abundant rhizobium (Barrett *et al.,* 2012). However, this process should minimize plant-rhizobia interactions for fitness *within* rhizobia species as well, inconsistent with the pervasive genotype-by-genotype interactions documented between *M. truncatula* and *E. meliloti* (Heath *et al.,* 2012).

### Complementarity of phenotypic and genotypic approaches

In the present study, we took advantage of a pre-existing genomic dataset to complement and extend our test for symbiont local adaptation using a classic reciprocal inoculation experiment. Our genomic outlier analysis also did not produce evidence of symbiont local adaptation, possibly because of our low sample size and low SNP coverage in our data set. However, we believe that combining an experimental approach and genomics is an innovative and powerful way to test for local adaptation that should be applied more broadly. Although genome scans and reciprocal inoculation experiments are typically treated as alternatives because they draw on fundamentally different data, together the two approaches constitute a rigorous test for local adaptation in environmentally sensitive symbioses such as the legume-rhizobia mutualism. Combined, the two approaches integrate over the effects of all loci in the genome (reciprocal inoculation experiments) and across ancillary environmental variation (genome scans), producing inferences that are less vulnerable to the weaknesses of either method (Buehler *et al.,* 2014; de Villemereuil *et al.,* 2015; Jensen *et al.,* 2016). Studies of (symbiont) local adaptation should consider pairing phenotypic and genomic approaches to validate their results with independent lines of evidence and exclude alternative interpretations of the data (de Villemereuil *et al.,* 2015; Jensen *et al.,* 2016). Future directions for our research could include repeating the genomic outlier test with a higher quality genomic data set to determine whether the reciprocal inoculation and genome scan produce concordant results on symbiont local adaptation.

## Acknowledgements

Our work is supported by Discovery Grants and graduate fellowships from NSERC Canada, and the EEB Postdoctoral Fellowship at the University of Toronto. We thank Bruce Hall and Andrew Petrie for greenhouse assistance, and Adriana Salcedo, Michelle Afkhami, and Rebecca Batstone for advice on experimental design and analysis. We are grateful for feedback from an anonymous Associate Editor and two anonymous reviewers, which strengthened the manuscript.

## Data accessibility

Sequence data will be uploaded to NCBI. VCF files and data from the reciprocal inoculation experiment will be available on Dryad. GPS coordinates of sampled plant and rhizobia populations are reported in Table S1.

## Supplementary Methods

### Appendix 1

#### Bioinformatics and SNP discovery in Ensifer

We aligned forward and reverse rhizobia reads to the reference genome of *E. meliloti* strain 1021 (Galibert *et al.,* 2001) (NCBI references chromosome AIL591688, plasmid a AE006469, plasmid b AL591985) and the *E. medicae* strain WSM419 (Reeve *et al.,* 2010) (NCBI references chromosome 150026743 plasmid b 150030273, plasmid a 150031715, accessory plasmid 150032810) using BWA (Li and Durbin, 2009) and Stampy (Lunter and Goodson, 2011) with default parameters and the bamkeepgoodreads parameter. We assigned bacterial species using a combination of the percentage of reads mapping to one reference genome, and sequences at the 16S rDNA locus (NCBI gene references 1234653 and 5324158, respectively), which differs between *E. medicae* and *E. meliloti* (Rome *et al.,* 1997). We used Integrative Genomics Viewer to visualize and check alignment quality (Robinson *et al.,* 2011). In general, 69.99 - 94.02% (median 84.71%) of reads per sample mapped to the *E. meliloti* reference genome, and 69.32 - 92.48% (median 83.49%) mapped to the *E. medicae* genome.

We used PICARD tools to format, sort, and remove duplicates in sequence alignments. We applied GATK version 3 indel realignment and GATK Unified Genotyper SNP discovery on all bacteria alignments (McKenna *et al.,* 2010) with ploidy set to haploid. We used the Select Variants parameter in GATK to select SNP variants only. We used standard hard filtering parameters and variant quality score recalibration on SNP discovery according to GATK Best Practices (DePristo *et al.,* 2011; Van der Auwera *et al.,* 2013). We filtered rhizobia SNPs for a minimum read depth (DP) of 20, a maximum DP of 226 for *E. meliloti* (230 for *E. medicae),* and a genotype quality (GP) of 30 using vcftools (Danecek *et al.,* 2011). We removed indels and sites with more than 10% of missing data from both *E. meliloti* and *E. medicae* data files. We identified synonymous SNPs using SnpEff (Cingolani, Platts, *et al.,* 2012b) and SnpSift (Cingolani, Patel, *et al.,* 2012a), using reference files GCA_000017145.1.22 and GCA_000006965.1.22 (for *E. medicae* and *E. meliloti,* respectively) in the pre-built database. We used the ANN annotation parameter in SnpSift to identify SNPs as synonymous variants and missense variants.

#### Bioinformatics and SNP discovery in Medicago

We called *Medicago* SNPs in GBS samples by following the three-stage pipeline in the program Stacks (Catchen *et al.,* 2011; 2013): cleaning raw data, building loci, and identifying SNPs. We trimmed reads to 64 bp and filtered reads by a phred score of 33, the default value for GSB reads sequenced on Illumina 2000/2500 machine. We built loci for *M. lupulina* using the *de novo* approach in Stacks (denovo_map command), setting the –m parameter at 5, the –M parameter at 1, and the -n parameter at 1. In the final stage of the pipeline, we identified SNPs under the populations command by setting the -m parameter at 5. We filtered SNPs by removing indels, removing sites with more than 10% of missing data, and removing sites that were less than 64 bps apart with vcftools (Danecek *et al.,* 2011). We also excluded 9 SNPs with heterozygosity that was higher than expected under Hardy-Weinberg.

### Appendix 2

#### Genomic outlier tests

We first performed the BLAST test in two ways: first using the range-wide sample of plants that hosted different bacterial species (73 plant individuals), and second, focusing on southern Ontario samples (49 plant individuals). We performed the latter test because of the possibility that many loci unrelated to bacterial specificity (e.g., climatic adaptation) could be differentiated between southern Ontario and the mid-Atlantic United States due to environmental gradients that covary with bacterial species composition.

Outlier loci detected in genotyping-by-sequencing (GBS) data are rarely the actual loci responsible for adaptation; instead, they are usually in linkage disequilibrium (LD) with the causal genes. To account for this possibility, we then searched for genes involved in the legume-rhizobia symbiosis within either 5 or 10 kb of the *M. truncatula* orthologs of the outlier loci that we detected in both the range-wide and Ontario samples. This approach assumes synteny between *M. truncatula* and *M. lupulina.* We chose 5 and 10 kb based on the scale of LD in *M. truncatula* (Branca *et al.,* 2011). While the scale of LD between even closely related species is likely to differ based on mutation rates, recombination, population structure, and a host of other demographic and evolutionary factors, we viewed this approach as superior to simply confining our searches to the GBS loci without accounting for their potential LD with causal genes.

Finally, we measured the distance between the *M. truncatula* orthologs of the outlier loci that we detected in both the range-wide and Ontario samples and key *M. truncatula* genes involved in the rhizobia symbiosis (again assuming synteny between *M. truncatula* and *M. lupulina).* We considered genes involved in the initial signal exchange between the legume and rhizobia (NSP, IPD3, and DMI1-DMI3); genes involved in infection thread development (LIN); and genes involved in both rhizobia signaling and infection (NFP, LYK3, and NIN) (Jones *et al.,* 2007; Oldroyd *et al.,* 2009; Young *et al.,* 2011; Oldroyd, 2013; Tang *et al.,* 2014).

### Appendix 3

#### Genomic outlier results

We identified a distribution of X^T^X statistics around the null expectation of X^T^X = 2, reflecting the 2 populations assigned in Bayenv2 (M. *lupulina* plants hosting *E. medicae* and plants hosting *E. meliloti).* In the range-wide sample, 16% (354 of 2209) of SNPs had X^T^X scores greater than the null expectation of 2; in the Ontario sample, 29% (573 of 1977) of SNPs had X^T^X scores greater than 2. We detected a range of alignment scores when we used BLAST to align outlier loci with top X^T^X statistics from the whole sample and the Ontario sample to the *M. truncatula* reference genome (Table 2). The loci mapped to several different chromosomes in the *M. truncatula* reference genome.

Of the top 1% of SNPs detected in the range-wide sample (20 SNPs total), eight were associated with a specific *M. truncatula* gene (BLAST scores: 35.6 – 102; E value: 1.00e-19 – 0.31). Higher BLAST scores reflect higher-quality alignments; these scores indicate that our sequences generally aligned moderately well to the *M. truncatula* genome. E (expectation-values reflect the number of hits expected by chance, so lower E-values indicate better matches. These 8 loci did not map to any genes known to be involved in the legume-rhizobia mutualism. The remaining 12 loci did not map to a specific gene in the *M. truncatula* genome (BLAST scores: 35.6 – 102; E-values: 3.00e-20 – 3.10e-1).

The results were qualitatively similar for the Ontario sample (Table 2). The BLAST scores of the top 1% of outlier SNPs (20 SNPs total) ranged from 37.4 to 111 (E-values: 5.00e-24 - 8.90e-2). Twelve of the top 1% of SNPs in the Ontario sample mapped to genes that are not known to be involved in the legume-rhizobia mutualism. The remaining eight loci did not associate with a specific gene in the *M. truncatula* annotated genome. The BLAST scores for these loci were similar to the twelve loci that did map to specific *M. truncatula* genes (score: 35.6 – 95.1; E-value: 3.00e-20 –3.10e-1).

There were only three outlier loci that appeared in the top 1% of SNPs in both the rangewide sample and the Ontario sample (Table 3). These loci mapped to chromosomes 1, 5, and 7 in the *M. truncatula* genome, but did not map to a specific gene. No genes found within 5 or 10kb of the *M. truncatula* orthologs of these three outliers are known to be involved in the legume-rhizobia symbiosis (assuming synteny between *M. truncatula* and *M. lupulina).* The *M. truncatula* ortholog of the outlier on chromosome 5 had two genes within 5 kb, a phosphate putative gene and a Ty3/Gypsy polyprotein/retrotransposon. The ortholog of the outlier on chromosome 1 had no genes within a 5 kb window, and the ortholog of the outlier on chromosome 7 had two genes within 5 kb, a DUF247 domain protein and a Gypsy-likepolyprotein/retrotransposon putative gene. When we increased our window size to 10 kb we found more genes, but none related to infection with rhizobia. For example, the *M. truncatula* ortholog of the outlier on chromosome 5 was close to a DUF679 domain membrane protein and an alpha/beta fold hydrolase putative gene. The ortholog of the outlier on chromosome 1 had a reverse transcriptase zinc binding protein and a homeobox knotted-like protein in its 10 kb window. The ortholog of the outlier on chromosome 7 had a phosphoenolpyruvate carboxylase within its 10 kb window, along with several putative proteins.

Finally, we calculated the distance in base pairs between the *M. truncatula* orthologs of the three outlier loci found in both the range-wide and Ontario analyses and several genes involved in *Medicago-rhizobia* association. None of the symbiosis genes that we considered were close to the orthologs of any of these three outliers. Most of the symbiosis genes are located on chromosome 5, but none were close to the ortholog of the outlier locus on chromosome 5 (Table 4). The ortholog of the outlier locus on chromosome 1 was approximately 35,822 kb away from the only symbiosis gene we considered that is located on chromosome 1 (LIN). The remaining two symbiosis genes—DMI1 and DMI2—are located on chromosomes 2 and 8 (Ané *et al.,* 2002), neither of which contained any outlier loci in our analysis.

**Table S1.**
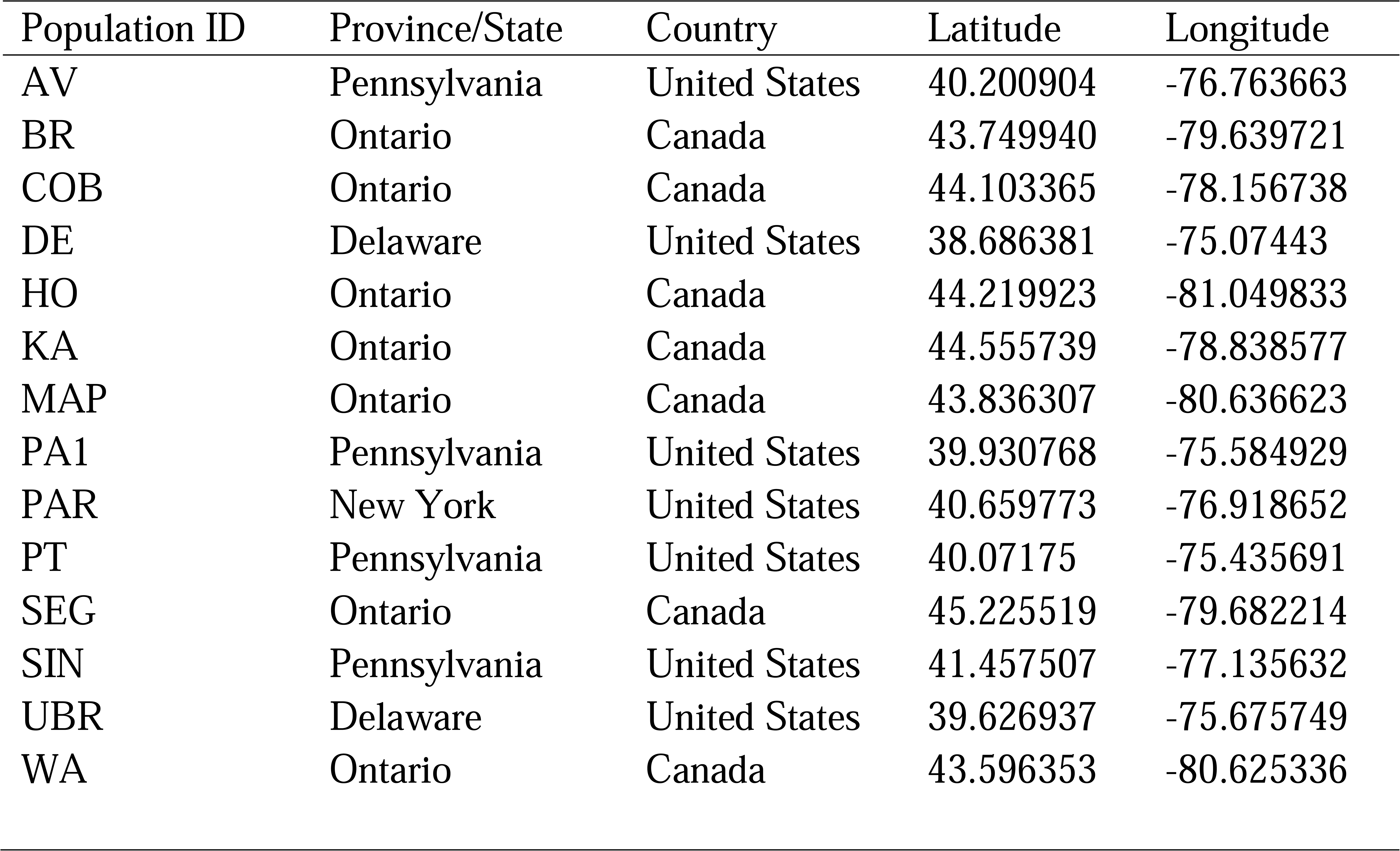
Locations of *M. lupulina* and *Ensifer* populations used in population genetic analysis.

**Table S2.** Summary statistics of Bayenv2 and BLAST results for the top 1% of SNPs in the X^T^X outlier analysis.

| Ontario sample |  |  |  |  |  |
| --- | --- | --- | --- | --- | --- |
| X<sup>T</sup>X | SNP identity | BLAST | Query cover | E value | Gene |
| 4.20 | 585117 | 37.4 | 56 | 0.089 | transmembrane protein |
| 3.83 | 1192907 | 93.3 | 100 | 1.00E-18 | indole-3-glycerol phosphate lyase IGL1 |
| 3.82 | 187811 | 93.3 | 98 | 1.00E-18 | toprim domain protein |
| 3.75 | 1110167 | 107 | 100 | 6.00E-23 | TLD-domain nuclear protein |
| 3.73 | 1610082 | 71.6 | 81 | 4.00E-12 | no result |
| 3.65 | 229813 | 35.6 | 60 | 3.10E-01 | no result |
| 3.59 | 1959186 | 107 | 100 | 6.00E-23 | TPR repeat protein |
| 3.56 | 129152 | 41 | 59 | 7.00E-03 | no result |
| 3.55 | 884266 | 89.7 | 100 | 2.00E-17 | no result |
| 3.44 | 616912 | 95.1 | 98 | 4.00E-19 | no result |
| 3.43 | 97240 | 111 | 100 | 5.00E-24 | transportin-1 protein |
| 3.39 | 666854 | 87.8 | 100 | 6.00E-17 | no result |
| 3.39 | 713735 | 96.9 | 100 | 1.00E-19 | indole-3-glycerol phosphate lyase IGL1 |
| 3.33 | 1294820 | 69.8 | 95 | 2.00E-11 | copia-like polyprotein/retrotransposon |
| 3.33 | 686219 | 69.8 | 100 | 2.00E-11 | no result |
| 3.33 | 1463455 | 91.5 | 100 | 5.00E-18 | single-stranded nucleic acid-binding protein R3H |
| 3.29 | 671122 | 37.4 | 85 | 8.90E-02 | cysteinyl-tRNA synthetase |
| 3.26 | 1071597 | 98.7 | 100 | 3.00E-20 | no result |
| 3.25 | 109965 | no result | no result | no result | no result |
| 3.25 | 734196 | 78.8 | 100 | 3.00E-14 | no result |
| Range-wide sample |  |  |  |  |  |
| 3.25 | 825707 | 96.9 | 100 | 1.00E-19 | FAD/NAD(P)-binding oxidoreductase family protein |
| 3.06 | 1131532 | 89.7 | 100 | 2.00E-17 | novel plant SNARE-like protein |
| 3.02 | 1482582 | 35.6 | 39 | 0.31 | ASCH domain protein |
| 2.99 | 405789 | 89.7 | 93 | 2.00E-17 | CAAX amino terminal protease family protein |
| 2.99 | 511074 | 48.2 | 96 | 5.00E-05 | no result |
| 2.97 | 175058 | 55.4 | 96 | 3.00E-07 | no result |
| 2.95 | 485278 | 91.5 | 93 | 5.00E-18 | no result |
| 2.94 | 254373 | 59 | 100 | 3.00E-08 | CRS1/YhbY (CRM) domain protein |
| 2.92 | 1090953 | 55.4 | 95 | 3.00E-07 | galactose oxidase |
| 2.90 | 921907 | 93.3 | 100 | 1.00E-18 | DUF223 domain protein |
| 2.86 | 1071597 | 98.7 | 100 | 3.00E-20 | no result |
| 2.82 | 870953 | 35.6 | 51 | 3.10E-01 |  |
| 2.81 | 1590352 | 102 | 92 | 3.00E-21 | no result |
| 2.75 | 686219 | 69.8 | 100 | 2.00E-11 | no result |
| 2.73 | 774291 | 69.8 | 95 | 2.00E-11 | no result |
| 2.71 | 313573 | 69.8 | 98 | 2.00E-11 | no result |
| 2.68 | 1342844 | 60.8 | 90 | 8.00E-09 |  |
| 2.67 | 1217147 | 89.7 | 100 | 2.00E-17 | novel plant SNARE-like protein |
| 2.64 | 616912 | 84.2 | 100 | 7.00E-16 | no result |
| 2.64 | 1057909 | 66.2 | 70 | 2.00E-10 | no result |

**Table S3.**
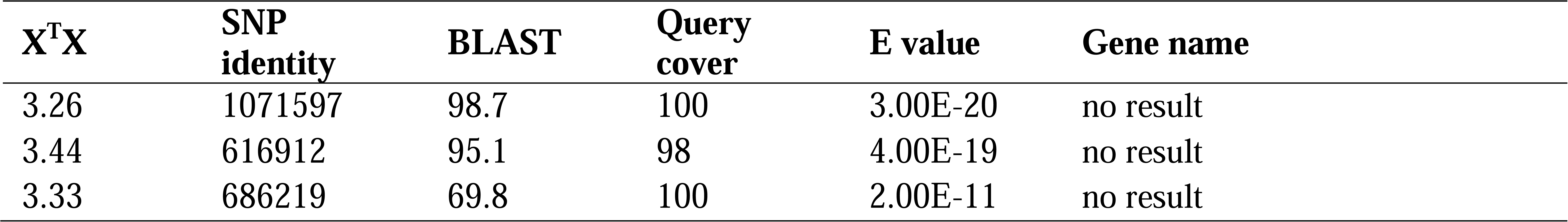
Outlier loci found in the top 1% of Bayenv2 results in both the *M. lupulina* range-wide and Ontario samples.

**Table S4.**
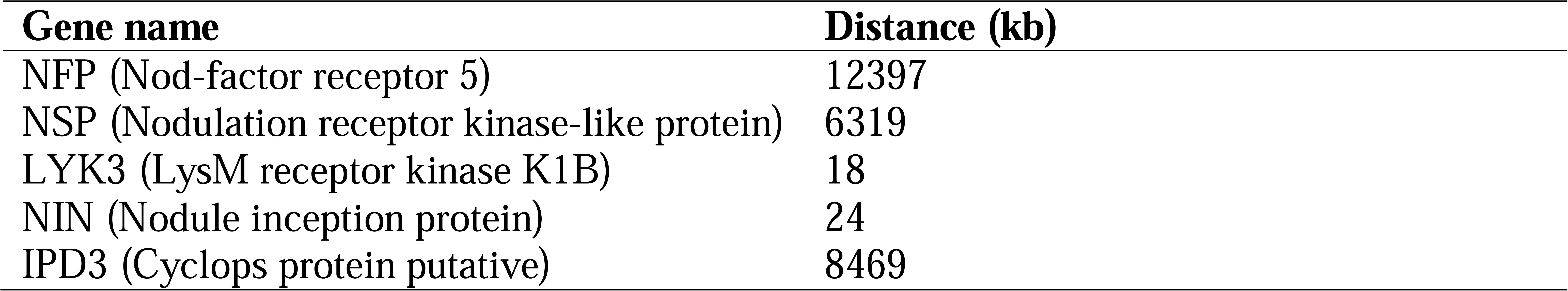
Base pair distances between the *M. truncatula* ortholog of the outlier locus on chromosome 5 and well-characterized nodulation and rhizobial infection genes in *M. truncatula.*

**Figure S1.**
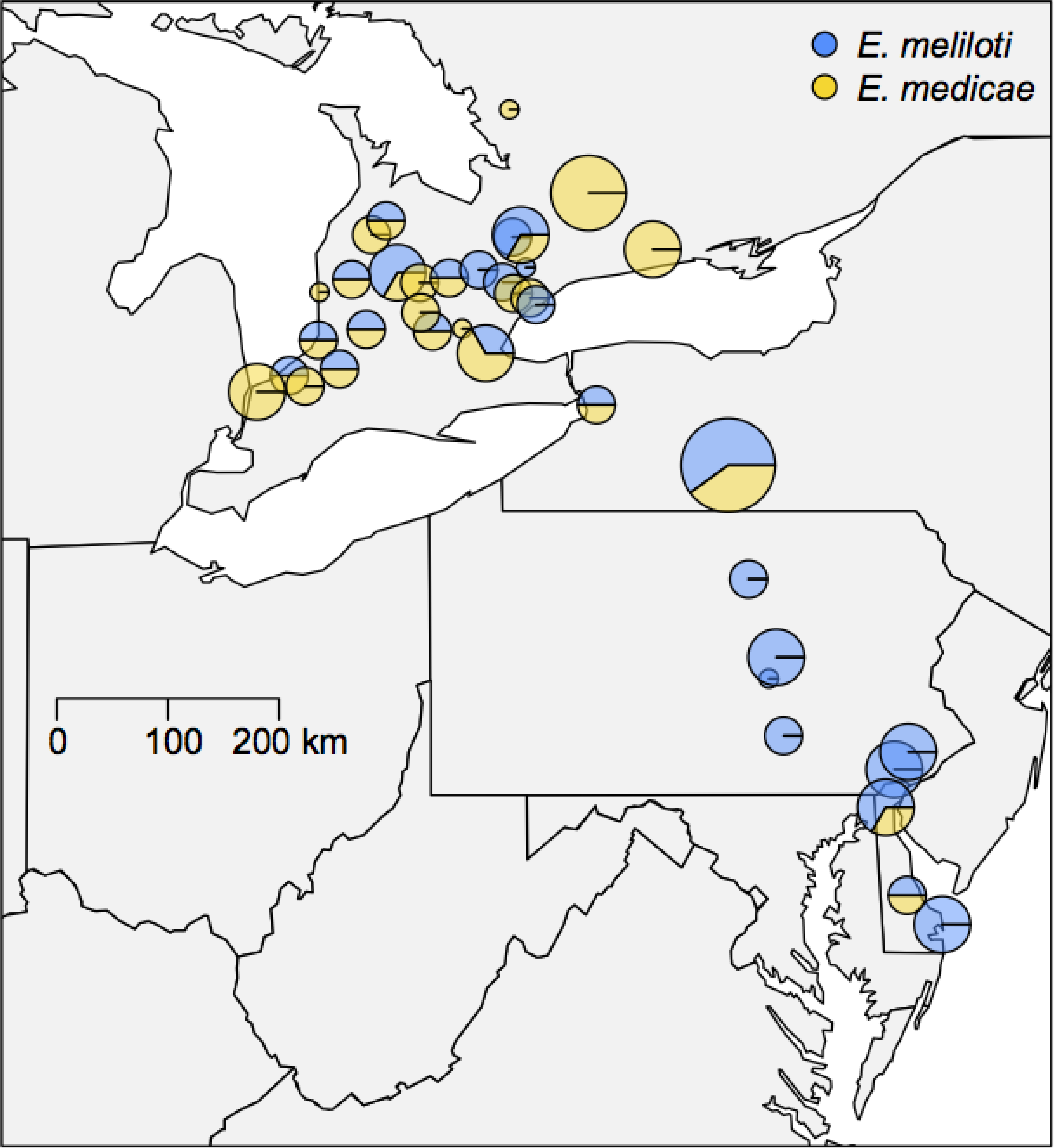
Locations and bacterial species present in all 39 sampled *M. lupulina* populations. The size of each circle corresponds to the number of sampled plant individuals for which their bacterial partner was identified to species. The colors correspond to the fraction of plants partnered with *E. meliloti* (blue) and *E. medicae* (yellow). The map is modified from Figure 2 in Harrison et al. (2017).

